# Dissociable components of the reward circuit are involved in appraisal versus choice

**DOI:** 10.1101/172320

**Authors:** Amitai Shenhav, Uma R. Karmarkar

## Abstract

People can evaluate a set of options as a whole, or they can approach those same options with the purpose of making a choice between them. A common network has been implicated across these two types of evaluations, including regions of ventromedial prefrontal cortex and the posterior midline. We test the hypothesis that sub-components of this reward circuit are differentially involved in triggering more automatic appraisal of one’s options (Dorsal Value Network) versus explicitly comparing between those options (Ventral Value Network). Participants undergoing fMRI were instructed to appraise how much they liked a set of products (Like) or to choose the product they most preferred (Choose). Activity in the Dorsal Value Network consistently tracked set liking, across both task-relevant (Like) and task-irrelevant (Choose) trials. In contrast, the Ventral Value Network was sensitive to evaluation condition (more active during Choose than Like trials). Within vmPFC, anatomically distinct regions were dissociated in their sensitivity to choice (ventrally, in medial OFC) versus appraisal (dorsally, in pregenual ACC). Dorsal regions additionally tracked decision certainty across both types of evaluation. These findings suggest that separable mechanisms drive decisions about how good one’s options are versus decisions about which option is best.

---

When people approach a display of options, they can evaluate them in at least two different ways. They may appraise the available options collectively as a set, such as when browsing a window display before walking into a store. Or they may focus on actively choosing between such options, indicating a preference for a favorite, such as when making a purchase. Though these modes of evaluation can involve identical stimuli, they seem to be phenomenologically distinguishable. Yet little is known about the degree to which they draw on shared or distinct mechanisms. Different lines of research have examined the neural mechanisms associated with appraising the value of an isolated item^1-3^ and others have examined the process of choosing one item from an assortment^4-6^. However, it remains unclear how people appraise a *set* of items when they don’t have the explicit goal of selecting between them. Put in the context of everyday decisions, to what extent are the mechanisms involved in browsing also involved in deciding what to buy?

Overlapping circuitry has been implicated in representing single-item preferences and multi-item choice^3-7^. This includes regions of ventral striatum, ventromedial prefrontal cortex (vmPFC), and the posterior midline. As a result, this is often treated as a single reward circuit, and mPFC in particular is often treated as one undifferentiated region implicated in the encoding and/or integration of reward. Indeed, across the literature, the label “ventromedial prefrontal cortex” is used in ways that cover a range of cortical subdivisions, including anatomically distinct regions around pregenual anterior cingulate cortex (pgACC; Area 24) and medial orbitofrontal cortex (mOFC; Area 14)^8-12^. However, some studies do suggest that finer distinctions can be made within this circuitry. Functional and structural differences have been documented between more dorsal and more ventral regions of vmPFC, and similarly for the posterior midline^5,8,13-16^. The same sets of regions have been dissociated using resting-state functional connectivity, between sub-networks of the Default Mode Network^17-20^.

Recent work by Shenhav and Buckner^17^ has provided indirect evidence suggesting that these separable sub-components of the reward network, and of medial prefrontal cortex in particular, may be differentially involved in appraisal versus choice evaluations of multi-item sets (Figure 2A). One of the networks identified by these studies consisted of mOFC, retrosplenial cortex (RSC), and left middle frontal gyrus (lMFG), which for brevity we will collectively refer to as a *Ventral Value Network*. This network was sensitive to a combination of choice difficulty and the value of one’s choice options. This pattern was consistent with prior evidence of mOFC’s involvement in value-based comparison in the service of a choice goal^21-24^.

**Figure 1.**
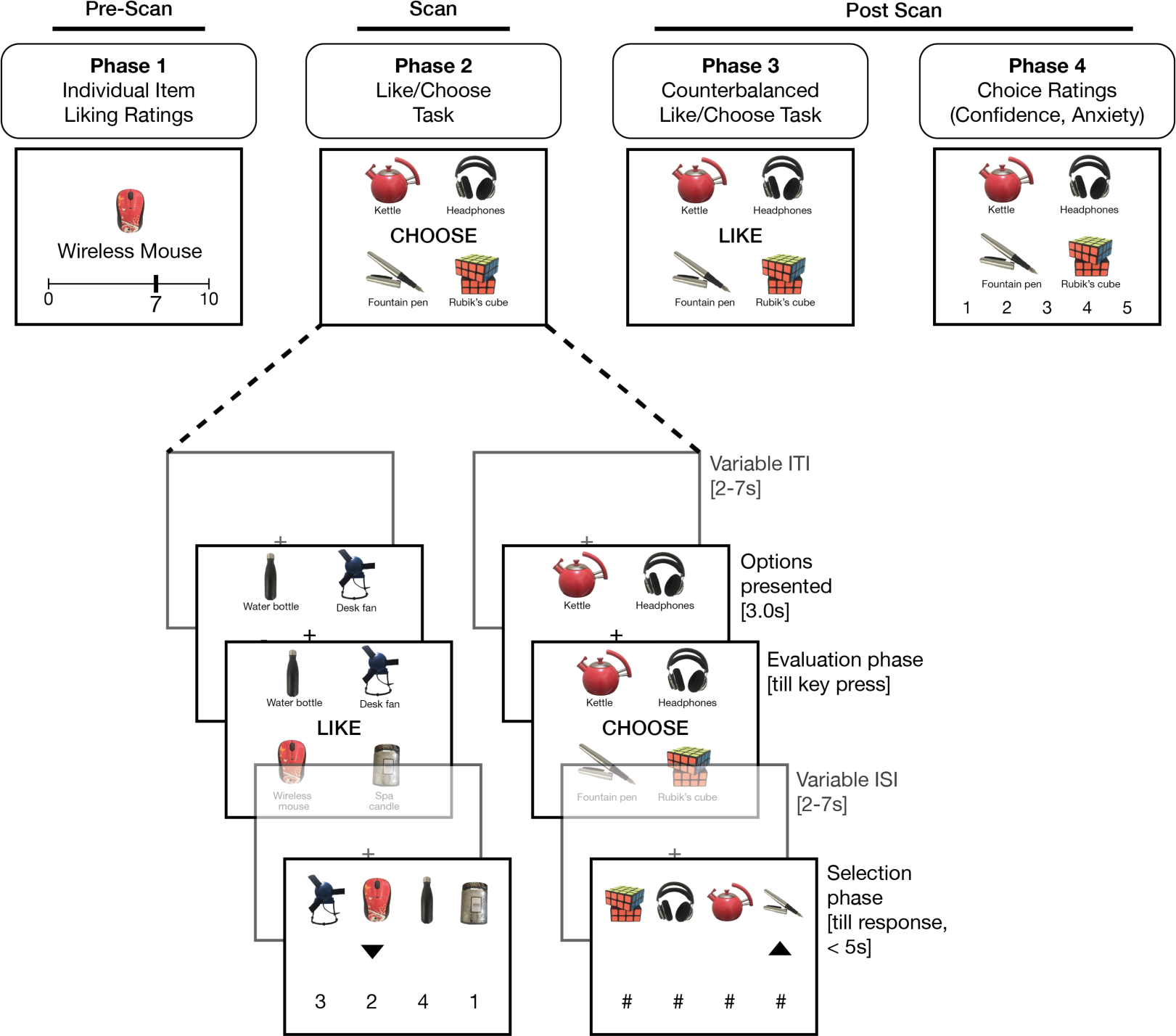
Participants rated each of the individual items (Phase 1) before performing the Like/Choose task (Phase 2). On each trial of Phase 2, participants viewed a set of four options and performed one of two kinds of evaluation. Options were presented for 3s at the start of each trial, and then one of two prompts appeared, indicating the task for that trial (CHOOSE or LIKE). Participants pressed a key once they completed their evaluation and, following a variable ISI, the options re-appeared along with numbers (Like) or # symbols (Choose). The participant used an arrow cursor to indicate their response, within a 5s deadline. Trials were followed by a variable ITI. In Phase 3, participants performed the counterbalanced evaluations for the Like/Choose task on each of the sets. In Phase 4, participants viewed each set again, and rated the confidence and anxiety experienced while choosing between the options. For copyright reasons, product images shown in the figure are representative of the study stimuli but not identical to the items used.

**Figure 2.**
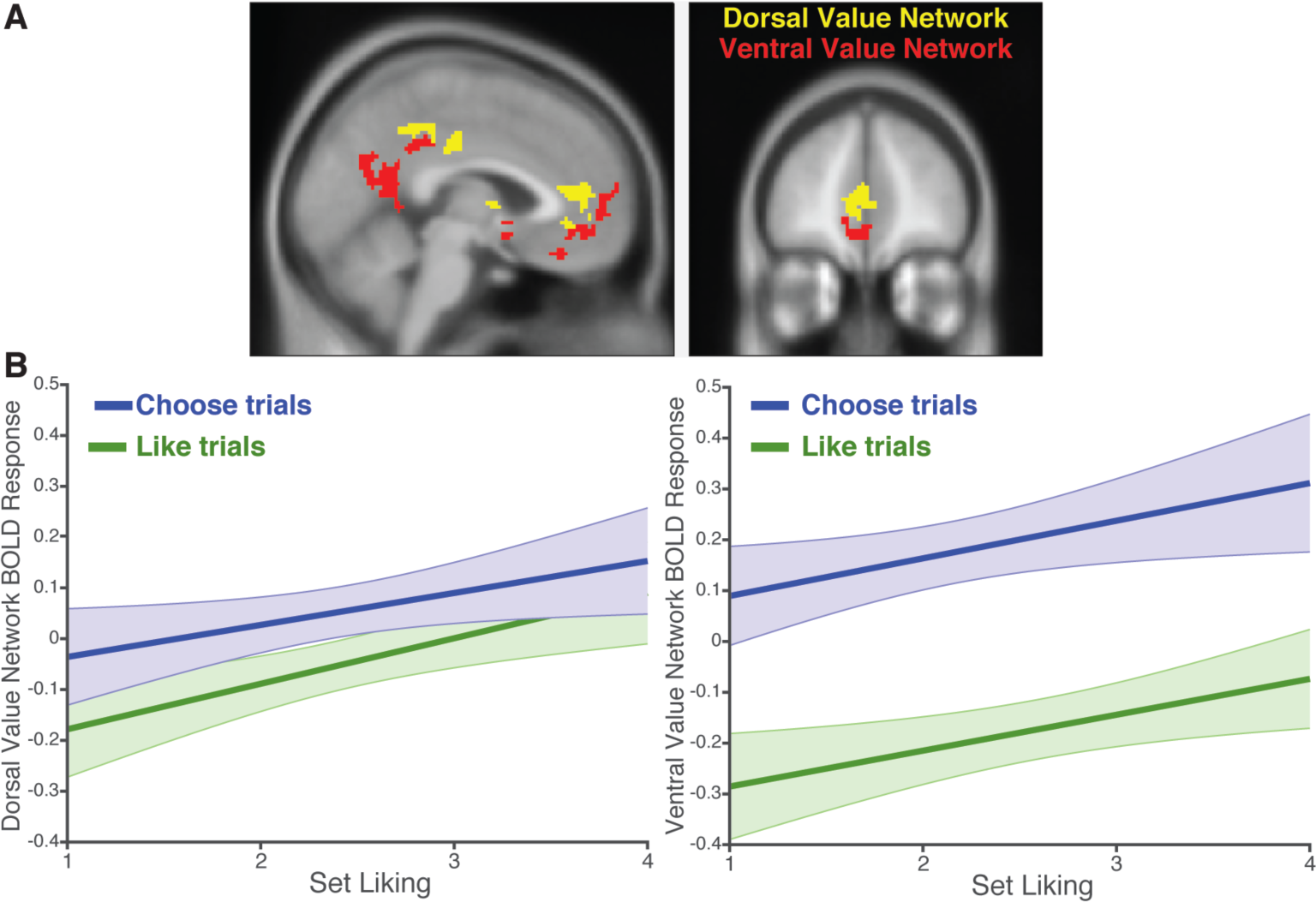
**A)** We defined ROIs for the Dorsal Value Network (yellow) and Ventral Value Network (red) based on regions that tracked choice-induced positive affect and difficult choice, respectively, in Shenhav & Buckner^17^. **B)** BOLD responses within the Dorsal Value Network (left) and Ventral Value Network (right) as set liking increases, as predicted by mixed-effects linear regressions. Both networks tracked set liking across both Like (green) and Choose (blue) conditions. However, the Ventral Value Network was significantly more sensitive to task condition than the Dorsal Value Network.

A second network – consisting of ventral striatum (VS), pgACC, and posterior cingulate cortex (PCC) – tracked the positive affect that was evoked by the choice options, but not the difficulty of choosing. Activity in these areas, which we will refer to as the *Dorsal Value Network,* was consistent with a more automatic or reflexive appraisal of these items rather than a process of comparing between them, paralleling findings from research on the automatic evaluation of liking – but not necessarily choice – of individual items^3,25-27^.

While prior research suggests a potential distinction between the kinds of evaluative goals that recruit different components of the reward circuit, this has yet to be directly tested using a task that varies these goals while holding other stimulus properties constant. Such a within-subject comparison is particularly critical given the wide between-experiment spatial heterogeneity of reward-related findings within vmPFC and other areas of these value networks^5-7,12^. Building on this, we examine whether activity in the Dorsal and Ventral Value Networks will diverge when people appraise a set of items versus when they select from among them. Given its proposed role in comparison-based choice, one possibility is that the Ventral Value Network (and mOFC in particular) will be more active when participants are making a choice relative to when they are appraising a set. In addition, given evidence for its role in reflexive evaluation of one’s options, we test whether the Dorsal Value Network will consistently signal the overall value of a set of items, regardless of whether this overall set value is task-relevant (appraisal) or task-irrelevant (choice).

To explore these questions we performed an fMRI study using a novel “Like vs. Choose” task (Figure 1). Participants viewed sets of products and either estimated how much they liked the whole set or selected the product they preferred most from it. Critically, both decisions required participants to consider the value of each item in the set, but differed in terms of whether they relied on a *composite* of those values or a *comparison* between them. For each product set, participants performed *either* a Like or Choose evaluation in the scanner. Afterwards, they performed the complementary evaluation outside of the scanner (i.e., rated their Liking for sets that they previously compared via Choose, and vice versa). We find that set liking (appraisal ratings) correlated best with the average value of the individual set items, and that the Dorsal Value Network signaled this liking irrespective of the task at hand. In contrast, the Ventral Value Network was particularly sensitive to whether the participants were choosing rather than appraising. We confirmed this dissociation within vmPFC, showing that pgACC was more sensitive to set liking and mOFC was more sensitive to evaluation type. Our task allowed for further comparisons between neural signals related to decision certainty for each type of evaluation. We show that both types of certainty are encoded in pgACC and the broader Dorsal Value Network, independently of set liking. Collectively, these findings point to separable mechanisms for browsing (appraisal) and choice, suggesting that the circuits that draw us to the store window may be different than those that guide our in-store purchases.

## Results

### Behavioral Results

#### Predictors of set liking

Our first goal was to understand how one’s appraisal of a set reflected the valued of the items in that set. For example the value of a set of options has been shown to reflect the average value of the individual items^28-30^. But it has also been shown that set value can be disproportionately influenced by the value of a preferred or salient item^31^ (cf. Ref. ^17^). We therefore considered whether the appraisal ratings provided during the Like task (*set liking*) were driven by the highest-valued item in the set, the average value of the items in the set (*average item value*), or some combination thereof. When regressing set liking on both of these variables, we found that it was most strongly associated with the average item value (β = 0.59, *t*(26.0) = 10.1, *p* < 0.001). The value of the best item in the set exerted a non-significant positive influence on set liking (*t*(25.6) = 1.7, *p* = 0.10). Notably, these results held when separately examining only the mixed-value item sets in which there was a greater difference between the highest item value (max) and the averaged item value than in the similar-value sets (average: β = 0.64, *t*(25.4) = 10.2, *p* < 0.001, max: β = 0.08, *t*(25.6) = 1.5, *p* = 0.16).

#### Distinct predictors of decision certainty during choice versus appraisal

We next examined the factors that contributed to how quickly people came to a Like or Choose decision in the Evaluation period (collapsing across trials completed inside and outside the scanner). These response times (RTs) offer a potential proxy for the strength of decision evidence provided on a given trial, and therefore the certainty with which that decision was made.

In the Like condition, evaluation RTs demonstrated a strong inverse U-shaped relationship with average item value, such that evaluations were fastest for the highest and lowest valued sets (linear effect: β = -0.09, *t*(24.6) = -3.4, *p* = 0.003, quadratic effect: β = -0.11, *t*(25.3) = -3.9, *p* < 0.001). We observed the same U-shaped pattern when regressing these RTs on set liking (linear effect: β = -0.03, *t*(24.6) = -1.2, *p* = 0.24, quadratic effect: β = -0.09, *t*(28.9) = -2.8, *p* < 0.01), consistent with previous findings of increased certainty when responding at the extreme of a scale^26^. Accordingly, Like RTs were significantly faster when participants indicated either the least or most liking for a set rather than moderate liking (i.e., they decreased with *rating extremity*; β = -0.21, *t*(24.6) = -3.6, *p* = 0.001).

In the Choose condition, evaluation RTs were best predicted by the absolute difference between the value of the chosen item and the average of the remaining items’ values (*value difference*; β = -0.25, *t*(29.5) = -9.4, *p* < 0.001), consistent with previous findings^32,33^. An alternate formulation of value difference^33^, based on the difference between the chosen and next-best item, demonstrated a similar but weaker correlation with RT (Supplementary Results 1). Unlike the quadratic effect observed in the Like condition, Choose evaluations only demonstrated a linear influence of average item value (faster with more valuable choice sets; β = -0.09, *t*(25.8) = -4.6, *p* < 0.001), over and above the primary effect of value difference on these evaluations^17,34^. Each of our two tasks thus offered independent and task-relevant indices of decision certainty via RT. For Choose trials, this was value difference, and for Like trials this was rating extremity.

Comparing the two tasks, we found that evaluations were slower on Choose relative to Like trials (β = 0.24, *t*(26.0) = 3.6, *p* = 0.001); to control for this, we include evaluation time as a covariate in all GLMs where these conditions are compared. Despite such differences in RT, participants did not find the Choose task to be more difficult than the Like task (average difficulty ratings: Choose = 4.00, Like = 4.15, paired *t*(26) = -0.37, *p* = 0.72).

#### Behavioral evidence against task spillover effects

Given the within-subject experimental design, we examined factors related to whether participants may have been influenced by the alternate task while performing a given evaluation (e.g., whether they were actively performing both appraisals and choices on all trials). As evidence against this possibility, we found that choice RTs (mean = 4.0s, median = 3.1s) were overall similar to, or shorter than, choice RTs in three previous behavioral studies (reported in Ref. ^35^) that did not interleave appraisals (mean = 4.9s, median = 3.8s). This suggests that choice RTs in the current study were unlikely to reflect a sequential combination of choice and appraisal. More importantly, we found that peoples’ appraisals of a set were not predicted by Choose-related variables such as value difference or the time taken for Choose evaluations (|*t*s| < 1.95, *p*s > 0.05). Together with additional dissociations between neural representations of decision certainty across these tasks (discussed below), this suggests that participants were focused on the specific evaluation indicated for a given trial.

We also tested whether participants complied with our instructions to make their choices or appraisals during the Evaluation phase of the trial rather than the Selection phase at the end of the trial. Consistent with this expectation, we found that RTs in the Selection phase were not significantly affected by average item value or rating extremity for either the Like or Choose task (|*t*s| < 1.85, *p*s > 0.05) and were only weakly affected by value difference on Choose trials (β = - 0.04, *t*(93.8) = -2.1, *p* = 0.038). This suggests that participants were performing the overall task as intended, by primarily making their decisions during the Evaluation phase, before advancing to Selection.

### fMRI Results

Our primary neuroimaging analyses of interest focused on two network ROIs defined based on patterns of activity from studies reported in Shenhav & Buckner^17^. The Dorsal Value Network (including pgACC, PCC, and ventral striatum) was defined as the set of regions that significantly tracked positive affective reactions to a choice set; the Ventral Value Network (including mOFC, RSC, and left MFG) was defined as regions that had been associated with difficult high-valued choices (see Figure 2A).

#### Task-independent encoding of set liking

First, we examined the hypothesis that areas in the Dorsal Value Network would track choice set appraisal. Consistent with this, we found that activity in this network increased parametrically with participants’ ratings during the Like task (β = 0.09, *t*(25.6) = 3.3, *p* = 0.003). Second, we examined whether the Dorsal Value Network would track set liking *irrespective* of the task being performed at the time (i.e., whether the participant was appraising the set or comparing the products with one another). Since participants gave both Like and Choose behavioral responses for each choice set (across Phases 2 and 3), we were able to test for correlates of set liking on Choose trials as well. As in the Like trials, activity in the Dorsal Value Network also increased parametrically with set liking when participants were engaged in the Choose task (β = 0.08, *t*(26.5) = 2.5, *p* = 0.020) (Figure 2B, left).

Overall, the Dorsal Value Network tracked set liking across all trials (*p* < 0.005), with no additional interaction between liking ratings and task condition (Choose vs. Like, *p*=0.61; Table 1). The model demonstrating this effect of set liking also included the evaluation RT, which did not significantly correlate with activity in the Dorsal Value Network (*p*=0.43). When simultaneously regressing Dorsal Value Network activity against set liking, average item value, and chosen item value, we found that set liking continued to be a significant predictor (β = 0.10, *t*(26.2) = 4.1, *p*<0.001) whereas the other two variables were not (|*t*s| < 1.85, *p*s > 0.05), suggesting that activity in this network was better accounted for by the subjective evaluation of the set as a whole than by combinations of item ratings. (For isolated effects of each predictor, see Table S2).

**Table 1.** Regression estimates for regressions predicting BOLD activity in the Dorsal and Ventral Value Networks based on simultaneous predictors for task condition (Choose > Like), set liking, condition x liking interaction, (log) evaluation time, and network (VVN > DVN).

| Outcome | Predictor | $\beta$ | SEM | t | p |
| --- | --- | --- | --- | --- | --- |
| **Dorsal Value Network** |  |  |  |  |  |
|  | Condition (Ch > Lk) | 0.11 | 0.04 | 3.06 | 0.003 |
|  | Set liking | 0.09 | 0.02 | 3.25 | 0.003 |
|  | Condition*liking | -0.02 | 0.04 | -0.52 | 0.61 |
|  | Evaluation time | -0.02 | 0.02 | -0.81 | 0.43 |
| **Ventral Value Network** |  |  |  |  |  |
|  | Condition (Ch > Lk) | 0.36 | 0.05 | 7.06 | <0.001 |
|  | Set liking | 0.07 | 0.03 | 2.59 | 0.015 |
|  | Condition*liking | 0.02 | 0.05 | 0.46 | 0.65 |
|  | Evaluation time | 0.07 | 0.02 | 3.57 | <0.001 |
| **Both Networks** |  |  |  |  |  |
|  | Network (VVN > DVN) | -0.12 | 0.04 | -3.35 | 0.001 |
|  | Condition (Ch > Lk) | 0.11 | 0.04 | 2.88 | 0.006 |
|  | Set liking | 0.09 | 0.03 | 2.86 | 0.008 |
|  | Condition*liking | -0.02 | 0.05 | -0.42 | 0.68 |
|  | Evaluation time | -0.02 | 0.02 | -0.78 | 0.44 |
|  | Network*condition | 0.25 | 0.06 | 4.49 | <0.001 |
|  | Network*liking | -0.02 | 0.03 | -0.69 | 0.49 |
|  | Network*condition*liking | 0.04 | 0.05 | 0.86 | 0.39 |
|  | Network*evaluation time | 0.08 | 0.03 | 3.24 | 0.001 |

The Ventral Value Network was also sensitive to set liking across all trials (p < 0.02), with no significant difference between Choose and Like trials (p = 0.49; Table 1; Figure 2B, right). However, given the close proximity of these networks and the apparent spatial separation between Like and Choose-related activity observed in our whole-brain analyses (Figure S1A), subsequent analyses probed the distinction between key regions within these networks more directly (see *Dissociable roles for pgACC and mOFC* section below).

#### Differentiation of choice versus appraisal

Even though both the Like and Choose tasks require participants to evaluate each of the items, we predicted that more ventral areas such as mOFC would be more active when engaged in the comparative evaluation needed for choice (cf. Refs. ^2,12,24^). Consistent with this, we found that Ventral Value Network activity was substantially greater for Choose relative to Like trials (β = 0.36, *p*<0.001; Table 1; Figure 2B, right). Though activity in the Dorsal Value Network was also sensitive to task condition (β = 0.11, *p* = 0.003), we observed a network by task interaction demonstrating that the relative difference in activity for Choose versus Like was significantly greater in the Ventral Value Network (β = 0.25, *p*<0.001) (Table 1).

These targeted ROI analyses focused on spatially dissociable network ROIs defined based on previous studies^17^. For completeness, we performed a confirmatory whole-brain analysis on the current data (Figure S1A). We observe similar dorsal/ventral distinctions between the regions sensitive to set liking and those sensitive to task condition (choice > appraisal; see also Table S1). Follow-up analyses on regions within the Ventral and Dorsal Value Networks showed that within the Dorsal Value Network, set liking was significantly correlated with activity in VS and pgACC (*ps* < 0.05) but not with activity in PCC (*p* = 0.47) (Table S3). Within the Ventral Value Network, task condition was significantly (and similarly) correlated with activity in mOFC, MFG, and RSC (*ps* < 0.05).

#### Overlapping task-dependent representations of decision certainty

Our analyses of response times suggested that there are distinguishable sources of decision certainty between the Choose and Like conditions. More certain decisions tend to be faster than less certain decisions^36-38^, and we found that Choose evaluations were fastest when the difference between chosen and unchosen values was greatest (cf. Refs. ^32,34,39^). In contrast, Like evaluations were fastest when the Like rating was at one of the extremes of the Likert scale (cf. Ref. ^26^). As such, the design of our task offered a unique opportunity to directly compare the neural activity evoked by these two forms of decision certainty.

We find that the Dorsal Value Network tracked both types of decision certainty. During the Choose task, activity in this network increased with our estimate of choice certainty (value difference; β = 0.10, *p* < 0.005). During the Like task, activity in this network increased with our estimate of appraisal certainty (appraisal rating extremity; β = 0.14, *p* < 0.05) (Table 2; see also Figure S1B). Each of these regression models controlled for evaluation time and set liking. This overlap helps draw together separate lines of research demonstrating that regions of the Dorsal Value Network track appraisal certainty when rating the pleasantness of a single item^26,40^ and track choice certainty when selecting among multiple potential rewards (e.g., Refs. ^34,41-44^; see also Ref. ^45^).

**Table 2.** The Dorsal Value Network tracks task-relevant decision certainty. When simultaneously regressing BOLD activity in this network on appraisal certainty and choice certainty (covarying set liking and evaluation time), appraisal and choice certainty are significant predictors on Like and Choose trials, respectively.

| **Outcome** | **Predictor** | **$\beta$** | **SEM** | **t** | **p** |
| --- | --- | --- | --- | --- | --- |
| **Dorsal Value Network (Like Trials)** |  |  |  |  |  |
|  | **Appraisal certainty** | **0.14** | **0.06** | **2.25** | **0.031** |
|  | Choice certainty | -0.02 | 0.03 | -0.89 | 0.38 |
|  | Set liking | 0.10 | 0.03 | 3.52 | 0.002 |
|  | Evaluation time | -0.03 | 0.03 | -0.89 | 0.38 |
| **Dorsal Value Network (Choose Trials)** |  |  |  |  |  |
|  | Appraisal certainty | 0.04 | 0.07 | 0.55 | 0.59 |
|  | **Choice certainty** | **0.10** | **0.03** | **3.44** | **0.001** |
|  | Set liking | 0.09 | 0.03 | 2.60 | 0.015 |
|  | Evaluation time | 0.02 | 0.03 | 0.70 | 0.49 |

While the neural correlates of decision certainty were similar across Like and Choose trials, it is important to note that the nature of the signals were task-dependent. Dorsal Value Network activity was not positively correlated with choice certainty in the Like condition^1^ (β =-0.02, *p* = 0.38), nor with appraisal certainty in the Choose condition (β =0.04, *p* = 0.59). Thus, neural activity related to certainty was tied to the particular evaluation being performed on that trial rather than more general correlates of appraisal and choice certainty (e.g., an automatic encoding of value difference). The behavioral and neural evidence of task specificity in certainty responses also provides further evidence that participants were engaging in distinct evaluations based on the goal of the trial.

Separate from the correlates of liking and certainty we observed across these two networks, we were unable to find additional signals in either network related to the value of the set, the value of the chosen item, or the (signed or unsigned) difference between chosen and unchosen values (Supplementary Results 2, Table S2). These analyses were also unable to identify signals in these networks related to the subjective confidence in one’s choice on a given trial (Supplementary Results 3), but we are cautious about interpreting this finding given that these confidence ratings were mostly at ceiling. While our task was not designed to discriminate between BOLD activity during initial option presentation versus during the presentation of the task cues (e.g. Evaluation), analyses locked to the onset of option presentation demonstrated similar patterns within both networks (Table S5).

#### Dissociable roles for pgACC and mOFC

The ROI results above suggest that the Dorsal and Ventral Value Networks are, to different degrees, both sensitive to task condition (Choose vs. Like). These analyses found that activity in the Ventral Value Network also tracked set liking across conditions (β = 0.07, *t*(30.2) = 2.6, *p* = 0.014), like the Dorsal Value Network. However, because these networks were defined by patterns of activation in a previous study, they lack anatomical specificity. Based on our hypothesized distinctions between more dorsal and ventral sections of the areas spanning what has been collectively considered ventromedial prefrontal cortex^5,9,12^, we performed a final set of analyses within ROIs defined based on anatomical boundaries specific to pgACC (Area 24) and mOFC (Area 14)^8,9^. In particular, these analyses tested which of these two (proximal) regions accounted for more of the variance arising from the three variables of interest: task condition (choice vs. appraisal), set liking, and decision certainty.

Similar to the findings above, both regions were separately associated with each of the three variables of interest (pgACC: *t*_*condition*_(58.7) = 4.2, *p*<0.001, *t*_*liking*_(32.1) = 3.2, *p*<0.005, *t*_*certainty*_(38.9) = 4.0, *p*<0.001; mOFC; *t*_*condition*_(81.8) = 5.2, *p*<0.001, *t*_*liking*_(26.5) = 2.0, *p*<0.06, *t*_*certainty*_(155.6) = 2.3, *p*<0.05). When including both regions within the same regression^3^,^46^,^47^ we find that these functional relationships dissociate, with mOFC sensitive to task condition but not set liking or decision certainty, and pgACC demonstrating the opposite profile (Figure 3, Table 3). Furthermore, we find a significant interaction between the BOLD sensitivity of the two regions (mOFC vs. pgACC) across the variable types (task vs. set liking vs. certainty). Specifically, paired comparisons of regression coefficients across participants demonstrate that mOFC is a stronger predictor of task condition than pgACC (Wilcoxon signed-rank test: *z* = 2.09, *p* < 0.05). They additionally find that mOFC is a stronger predictor of task condition than it is of set liking or decision certainty (*zs* = 4.54, *ps* < 0.0001). Finally, these comparisons show that pgACC is a stronger predictor than mOFC of set liking (*z* = 2.79, *p* < 0.01) and decision certainty (*z* = 3.80, *p* < 0.0005).

**Table 3.** Regression estimates for regressions predicting set liking, decision certainty, and task condition based on simultaneous predictors for pgACC activity, mOFC activity, and (log) evaluation time.

| Outcome | Predictor | $\beta$ | SEM | t | p |
| --- | --- | --- | --- | --- | --- |
| Set liking |  |  |  |  |  |
|  | pgACC | 0.06 | 0.02 | 2.56 | 0.016 |
|  | mOFC | 0.02 | 0.02 | 0.70 | 0.49 |
|  | Evaluation time | -0.05 | 0.02 | -1.99 | 0.057 |
| **Decision certainty** |  |  |  |  |  |
|  | **pgACC** | **0.06** | **0.02** | **3.10** | **<0.005** |
|  | mOFC | -0.01 | 0.02 | -0.74 | 0.46 |
|  | Evaluation time | -0.22 | 0.03 | -7.14 | <0.001 |
| **Task condition** |  |  |  |  |  |
|  | pgACC | 0.07 | 0.05 | 1.54 | 0.12 |
|  | **mOFC** | **0.16** | **0.05** | **3.14** | **<0.005** |
|  | Evaluation time | 0.26 | 0.08 | 3.50 | <0.005 |

**Figure 3.**
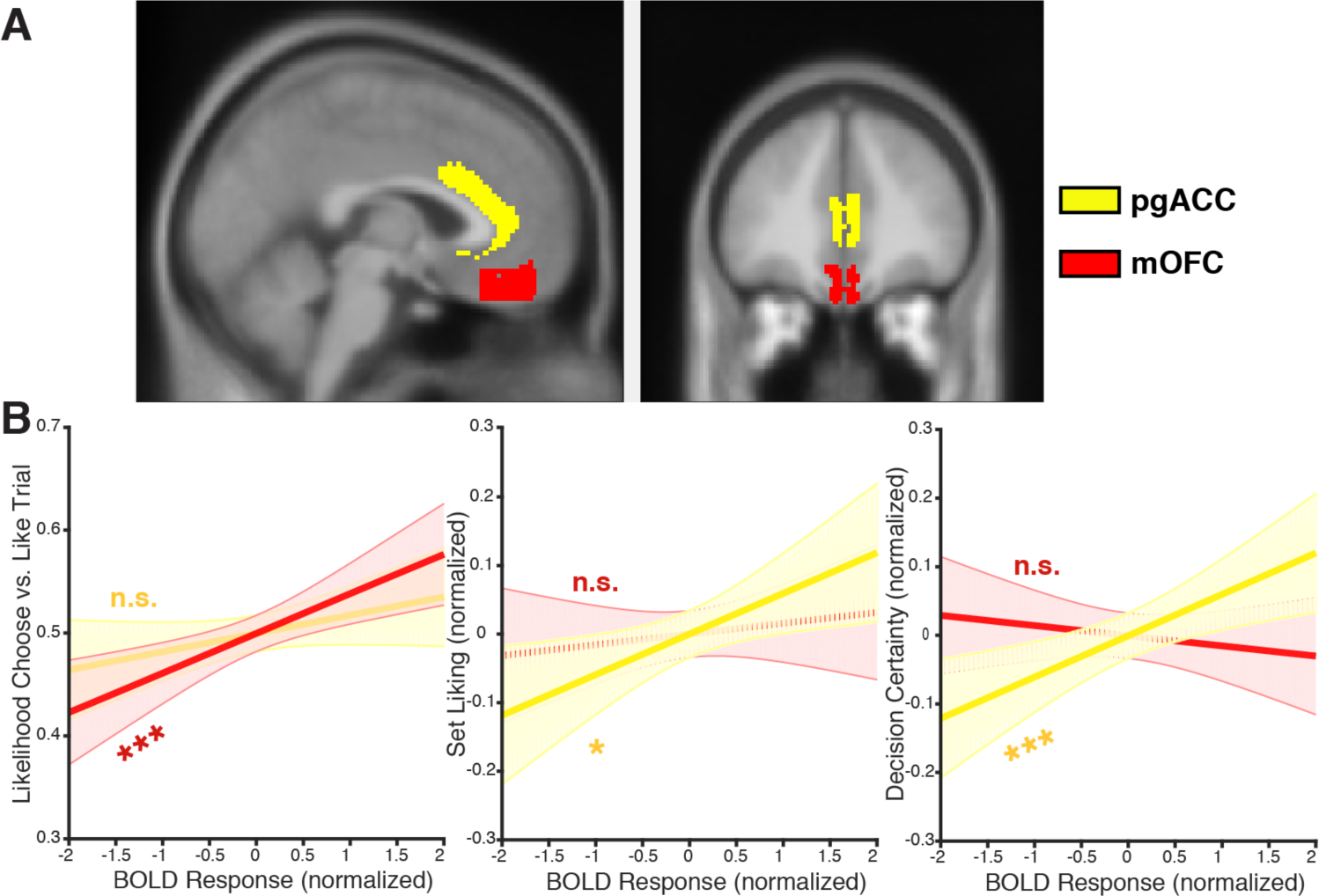
**A)** We defined ROIs for pgACC (yellow) and mOFC (red) based on a probabilistic anatomic atlas by Córcoles-Parada and colleagues^8^, thresholded at a minimum 50% probability of a given voxel being classified as part of a given anatomical region. **B)** BOLD estimates from separate regressions predicting task condition (**left**), set liking (**middle**), and decision certainty (**right**) based on activity in both pgACC and mOFC. When controlling for activity in pgACC, mOFC is significantly associated with task condition but not liking or certainty. Conversely, when controlling for activity in mOFC, pgACC is significantly associated with liking and certainty but not task condition. \**p*<0.05 \*\*\**p*<0.005

## Discussion

People can appraise potential rewards with or without the intent to choose between them. To better understand the mechanisms underlying these different but related processes, we directly contrasted appraisal and choice and uncovered two key findings. First, a Ventral Value Network – comprising regions of mOFC, RSC, and MFG – was more active when participants compared options to make a choice, rather than when they appraised the overall value of the set. Second, a Dorsal Value Network – comprising regions of pgACC, PCC, and ventral striatum – tracked how much participants liked those options, irrespective of whether they were tasked with reporting set liking on a given trial. The Dorsal Value Network also tracked the individual’s certainty in the evaluation they were making on that trial. Together these results provide valuable insight into the mechanisms underlying different forms (or components) of reward evaluation.

A substantial body of work has reported value signals across regions of the networks we explored, both when the task requires participants to make a choice^4-6,12^ and when it does not^1,48-50^. A common interpretation of these findings is that the value signals being uncovered in the two cases reflect a common valuation circuit engaged in an implicit decision process, irrespective of the task being performed. However, our findings speak to an alternative interpretation^12,17^, which proposes that these signals reflect two different valuation processes: one involving the triggering of affective (cf. Pavlovian) associations, the other involving a direct comparison between those outcomes, potentially via mutual inhibition^23,34,51^.

In line with the latter proposal, our results suggest that regions such as pgACC, ventral striatum and connected regions may signal reflexive affective associations while mOFC and connected regions may be more directly involved in active choice comparison. We did find evidence that both networks were sensitive to set liking and task condition, consistent with the fact that these networks included adjacent sets of voxels. However, when specifically examining subdivisions of vmPFC (pgACC and mOFC) as simultaneous predictors, we find that they demonstrate dissociable roles in signaling set liking versus choice comparison. These findings leave open the possibility that both networks carry signals related to task condition and set liking, and/or that they reflect different stages of evaluation. Future research using measures with higher temporal resolution could potentially shed light on this issue by examining the relative timing of these neural signals during appraisal and choice.

While only one of our conditions required participants to compare their options, both conditions required participants to select a response, whether it was selecting one of the four items or selecting their appraisal (liking) rating. Regions of our networks of interest have been shown to track variables related to choice selection, for instance the relative value of the chosen versus the unchosen item(s)^39,52^. It has been proposed that these value difference signals may reflect the decision output itself^34,53^, and/or that they reflect a metacognitive signal related to the confidence or ease with which the decision was made^41,54^. Separate research has shown that some of these same regions track similar metacognitive signals when rating one’s liking of an individual item^26^ – these studies show that pgACC tracks one’s confidence in that rating, indexed by how close the rating was to the endpoint of the scale (see also Refs. ^40,55^). We replicated such results in the current study, showing that the Dorsal Value Network tracked value difference signals during Choose trials and the extremity of ratings during Like trials. Importantly, because participants did not have information about the actions required to submit their choice when they were evaluating their options, neither these findings nor the findings above can be attributed to valuation/selection of specific motor effectors.

Our certainty-related results are notable both for the dissociation and for the integration they reveal. For example, in considering the task, one might ask whether participants automatically engage in both appraisal and choice during the evaluation period irrespective of task instruction, and then “gate” the relevant response during the selection period. If that were true, we would expect to see both appraisal- and choice-related decision certainty signaled on each trial. Instead, we found that certainty signals in the Dorsal Value Network only reflected the relevant task and not the irrelevant task (i.e., appraisal certainty only on Like trials, choice certainty only on Choose trials), suggesting that participants were only generating a response for a single task on each trial. This dissociation is particularly striking when juxtaposed with our finding that these regions tracked set liking in a task-*independent* manner, on both Like and Choose trials.

The fact that set liking and decision certainty signals converge in pgACC is also noteworthy (cf. Refs. ^26,40^). There are at least two intriguing explanations for these convergent signals. One possibility is that liking and certainty signals *both* reflect forms of affective appraisal rather than an explicit component of a decision process. Under this account, the Dorsal Value Network reflects one’s affective state, which is increasingly positive when viewing good options (irrespective of the task) and when more confident in one’s decision. A second explanation follows from models of choice comparison predicting that a region involved in such comparison should encode the overall value of a choice set (the key predictor of set liking in our study) and the relative value of the chosen item (the key predictor of certainty for our Choose trials)^34,56^. However, this prediction has only been applied to choice (not appraisal), and has typically been proposed for mOFC (not pgACC) because mOFC is believed to be more involved in choice comparison.

Indeed, we did find that mOFC activity was greater for choice than appraisal trials, consistent with the choice comparison account. However, after controlling for pgACC activity, we did not find additional choice-relevant value signals in mOFC (e.g., average item value, value difference). The absence of these independent value-related signals deserves further investigation. For example, it may reflect a non-linear relationship between activity in this region and the inhibitory dynamics of value-based choice. However, it is also possible that the increases in mOFC activity we observed for the Choose relative to the Like task reflected processes that contribute to the comparison process in other ways. For instance, compared to appraisal, choice may require additional generation of associated items and contexts in episodic memory, and these processes have been shown to be associated with mOFC and its broader network^18,57,58^.

By establishing a direct comparison between the processes involved in appraisal and choice, the current study can also be combined with other findings to offer valuable insight into the processes that drive us to prefer one option set over another. Indeed, this work suggests a tension between networks that support opposing preferences. While activity in the Dorsal Value Network and associated positive feelings may scale with the rewards on offer, separate networks appear to be involved in managing the decision process and in signaling the subjective costs (e.g., anxiety) of overcoming conflict between salient options^17,35^. As a result, whether one is appraising a set or choosing from it can affect how demanding or aversive their evaluation will be^35^. Improving our understanding of the dissociations between these circuits may therefore hold promise for reducing the costliness of transitioning from being a browser to being a chooser.

Our findings also have potential implications for research into reward-related impulsivity and its relationship to other forms of valuation. They suggest that the systems that signal the availability of reward (and potentially propel approach behavior towards those rewards) are at least partially dissociable from the circuits involved in selecting among those rewards. This offers a potential mechanism for divergent phenomenology when “browsing” a display or store window versus when “buying” or selecting from the options available inside the store. In addition, the patterns of findings we observed in the Dorsal Value Network have intriguing parallels in research on reward reactivity in impulse control disorders (e.g., to food and drug cues^59,60^), and are consistent with the possibility that these rewards can drive reward-seeking behavior independently of goal-directed comparison^61,62^. Additional research on this topic and its expression in real world contexts could thus benefit by extending to approach-related behavior linking appraisal and choice (e.g., the act of entering the store).

## Methods

### Participants

Thirty-one individuals (54.8% female, M_age_ = 25.0, SD_age_ = 4.4) completed this study. Of these, one was excluded from our analyses for excessive head motion (between-volume rotational movement two orders of magnitude greater than the group SD) and three were excluded due to an error in recording behavioral responses on the Like/Choose Task, leaving 27 individuals (55.6% female, M_age_ = 24.2, SD_age_ = 4.0) in the analyzed sample. Three other individuals initiated sessions but did not get scanned or complete the task, one due to technical difficulties and the other two due to insufficient variability in their initial product ratings (preventing our automated algorithm from generating the necessary quantity of option sets). Participants who engaged in part or all of the task received monetary compensation for their time.

This study was approved by the Institutional Review Board at Harvard University. All subjects gave their written informed consent and all experiments were performed in accordance with relevant guidelines and regulations.

### Experimental Design

Participants performed the study in three phases occurring sequentially within the same experimental session. In Phase 1, participants evaluated how much they would hypothetically like to have each of a series of products on a scale of 0 (‘not at all’) to 10 (‘a great deal’). The product sets were partially tailored in relation to the participant’s gender (total products evaluated: males = 310, females = 328).

In Phase 2, participants performed the *Like/Choose Task* (LCT) while in the scanner (Figure 1). On each trial of this task, participants were shown four of the previously-rated products and asked to either evaluate the set as a whole (Like trials; 1-4 Likert scale ranging from lowest to highest set attractiveness) or select the product they most prefer (Choose trials). In order to ensure that participants were able to view all of the products displayed before the explicit evaluation period, the products appeared for 3s at the start of each trial without information about the type of evaluation required. A LIKE or CHOOSE cue then appeared on the screen to indicate the task on the current trial. In order to separate evaluation and response selection periods, after the cue appeared participants were given an unlimited amount of time to make their decision and instructed to press a key once they had made their decision. This keypress ended the *Evaluation* period and a fixation cross was shown on the screen for a variable inter-trial interval (ITI; 2-7s), followed by the *Selection* period.

For Like trials, the Selection period consisted of the numbers 1-4 appearing at the bottom of the screen and the four products appearing at the top of the screen. Both numbers and products were shown in a random horizontal arrangement. Participants used the keypad to move a cursor left (right index finger) or right (right ring finger) before submitting their response (right middle finger). Participants were required to indicate a response within 5s during the Selection period. This deadline was intended to reinforce the idea that the response should have already been determined during the Evaluation period. Subsequent RT analyses confirmed that participants were conforming to these expectations (see Behavioral Results). The Selection period for Choose trials took place in a nearly identical manner, except that the liking rating numbers were replaced by # symbols, and participants moved the cursor to indicate the item they wished to choose (See Figure 1). Following the Selection period, there was a 2-7s ITI prior to the start of the next trial. There were a total of 120 trials in the LCT task, broken up into two blocks of 60 trials each. Each block began with a 10s fixation period (to allow for T1 equilibration) and ended with a 10s fixation period. Given the free response nature of the trials, block lengths varied within and across participants based on the decision times within the corresponding block (mean: 17.7 minutes, SD: 1.66 minutes).

Given that the study purpose was to directly compare the evaluative processes involved in appraisal and choice prior to a final decision, the Like-Choose Task was not incentivized. This approach avoids confounds arising from potential subjective differences in the subjective value of the decisions’ outcomes. To encourage participants to attend to the items, they were informed that regardless of their task responses, they would have the opportunity to receive an item randomly selected from the products viewed (or a cash bonus) at the end of the experiment. The alternative cash bonus was equal to half the price of the randomly selected product.

Each of the 120 LCT trials featured a product set generated based on the participants’ own preferences. Briefly, products were rank-ordered on the basis of Phase 1 ratings, and the resulting distribution split into tertiles. In order to ensure sufficient variability in the overall and relative values of the product sets, we generated and combined two types of sets. *Similar-value* product sets were generated by selecting 60 non-overlapping sequences of four consecutively rank-ordered products (20 sets from each tertile). *Mixed-value* sets were generated by randomly sampling four products from across the entire value distribution (without replacement). Sets were constructed such that each product would appear exactly twice (once in a similar-value set, once in a mixed-value set). Each product set was only seen once while in the scanner, either in the Like or the Choose condition.

After exiting the scanner, in Phase 3, participants completed a counterbalanced version of the LCT (i.e. providing Like ratings for sets that had been presented in the Choose condition in the scanner, and vice versa). This counterbalanced task was otherwise identical to the in-scanner LCT except that the additional time at block start and end, pre-Evaluation option presentation time, and ITIs were all fixed at 1s. In Phase 4, participants rated their anxiety and confidence associated with each choice on a 5-point scale. (The anxiety ratings were taken in service of a separate set of hypotheses^35^, and are not reported further here.) At the very end of the session, participants were also asked to separately rate how difficult the Choose and Like tasks had felt on a 9-point scale.

### Neuroimaging Parameters

Scans were acquired on a Siemens Trio 3T scanner with a 12-channel phase-arrayed head coil, using the following gradient-echo planar imaging (EPI) sequence parameters: repetition time (TR) = 2500 ms; echo time (TE) = 30 ms; flip angle (FA) = 90°; 2.5mm voxels; 0.50 mm gap between slices; field of view (FOV): 210 x 210; interleaved acquisition; 37 slices. To reduce signal dropout in regions of interest, we used a rotated slice prescription (30° relative to AC/PC) and modified z-shim prepulse sequence. The slice prescription encompassed all ventral cortical structures but limited regions of dorsal posterior parietal cortex. Structural data were collected with T1-weighted multi-echo magnetization prepared rapid acquisition gradient echo image (MEMPRAGE) sequences using the following parameters: TR = 2200 ms; TE = 1.54 ms; FA = 7°; 1.2 mm isotropic voxels; FOV = 192 × 192. Head motion was restricted with a pillow and padded head clamps. Stimuli were generated using Matlab’s Psychophysics Toolbox and were viewed through a mirror mounted on the head coil. Participants used their right hand to respond with an MR-safe response keypad.

### Statistical Analysis: Behavioral Data

Behavioral data were analyzed with mixed-effects regressions, accounting for individual subject variance as random effects. Evaluation times were positively skewed and so were log-transformed before being analyzed. Set liking was treated as a continuous variable. We analyzed decision certainty by using indices previously validated for estimating certainty (or overall strength of evidence) for the respective form of evaluation. For Choose trials, certainty was estimated as the absolute difference between the value of the chosen item and the average of the remaining items (based on Phase 1 ratings) (cf. Refs. ^32,33^). For Like trials, it was estimated based on the extremity of the participant’s Like rating on a given trial, with a binary variable classifying responses of 1 or 4 (least or greatest liking) as high certainty and 2 or 3 (intermediate liking) as low certainty (cf. Ref. ^26^). We further analyzed how appraisal and evaluation time varied with the average value of the items in the set. Given that the number of items is held constant across trials, this study was not designed to distinguish between effects related to this average value estimate and those related to total set value.

### Statistical Analysis: Neuroimaging Data

#### Preprocessing

fMRI data were analyzed using SPM8 (Wellcome Department of Imaging Neuroscience, Institute of Neurology, London, UK). Preprocessing consisted of realigning volumes within participant, resampling to 2mm isotropic voxels, nonlinear transformation to align with a canonical T2 template, and spatial smoothing with a 6mm full-width at half-max (FWHM) Gaussian kernel.

#### Trial-wise ROI analyses

Preprocessed data were submitted to linear mixed-effects analyses, using a two-step procedure. First, we generated BOLD signal change estimates for each trial using a first-level general linear model (GLM) in SPM. This GLM separately modeled stick functions with onsets at the Evaluation and Selection period of each trial. Trials were concatenated across the two task blocks, and additional regressors were included to model within-block means and linear trends. Finally, in order to remove noise related to head motion and other potential nuisance variables. the GLM was estimated using a reweighted least squares approach (RobustWLS Toolbox^63^). After estimating this first-level GLM, we extracted beta estimates for each trial from our primary regions of interest (ROIs; see below), transformed these beta estimates with the hyperbolic arcsine function (to achieve normality), and then analyzed trial-to-trial variability in these BOLD estimates with linear mixed-effects regressions (using Matlab’s *fitlme* and *fitglme* functions). Fixed effect degrees of freedom were estimated using Satterthwaite approximation. Given that evaluation time varied across conditions, all regressions covaried this variable. Projected values (Ŷ) and confidence intervals shown in Figures 2 and 3 were generated using Matlab’s *predict* function, based on the relevant mixed-effects regressions. To compare beta coefficients associated with different ROIs across different types of regressions (*Dissociable roles for pgACC and mOFC* section), we extracted random effects coefficients from a version of the relevant mixed-effects regressions that omitted fixed effects of those variables. We then performed nonparametric paired comparisons on the random effects from the relevant regressions.

#### Regions of interest

In order to examine activity in the Dorsal and Ventral Value Networks (Figure 2A), we generated ROIs based on whole-brain statistical maps from two fMRI experiments reported in Shenhav & Buckner^17^. The Dorsal Value Network mask was defined based on regions in which activity had been correlated with positive affect experienced when viewing one’s options (pgACC, ventral striatum, and PCC). The Ventral Value Network was defined based on regions that had shown greater activity for choices between two high-value options versus choices between a high-value option and a low-value option (mOFC, RSC, left MFG). This contrast compared choice sets that were matched for the best outcome but differed in the difficulty of comparison. For each of these two contrasts, we generated a mask consisting of a conjunction of the network regions that were consistently active across both previous fMRI studies (based on a voxelwise threshold of p<0.001 and cluster extent threshold of 50 voxels within each study). We excluded from each mask any voxels that intersected the Dorsal and Ventral Value networks, as well as any voxels that were part of a third network that tracked choice anxiety across these earlier studies (consisting primarily of dorsal ACC and anterior insula). Orthogonal analyses of this anxiety-related network are reported elsewhere^35^.

We followed up these functionally-defined network analyses with analyses that targeted anatomically-defined ROIs for pgACC (Area 24) and mOFC (Area 14) based on a probabilistic anatomic atlas by Córcoles-Parada and colleagues^8^ (Figure 4A). These ROIs were generated by thresholding the atlas such that each ROI only contained voxels with 50% or greater probability being classified as part of the given anatomical region.

## Data availability

All the data that support the reported findings are available from the authors upon reasonable request.

## Supporting information

## Acknowledgments

The authors are grateful to Elizabeth Beam, Marina Burke, Erin Guty, Tatiana Lau, Emily Levin and Erik Nook for assistance in data collection and Randy Buckner for helpful discussions. This work was funded by a postdoctoral fellowship from the CV Starr Foundation and a Center of Biomedical Research Excellence grant P20GM103645 from the National Institute of General Medical Sciences (A.S.).

## Author contributions

A.S. and U.K. designed the study and collected the data; A.S. analyzed the data; A.S. and U.K. wrote the manuscript.

## Competing interests

The authors declare no competing interests.

## Footnotes

1 The task-specificity of these value difference correlations also held when examining *signed* value difference (Chosen minus Unchosen) rather than the unsigned measure used as a proxy for choice certainty throughout the paper (Table S2).

