## Supplementary material for "Dissociable components of the reward circuit are involved in appraisal versus choice"

### Supplementary Methods

*1. Confirmatory whole-brain analyses.*

We supplemented our ROI analyses with whole-brain GLMs. For these analyses, first-level GLMs again separately modeled events occurring at Evaluation and Selection periods of each trial. The Evaluation period was modeled as a single event, modulated by a parameter of interest, including (log) evaluation time, task condition (indicator variable for Choose vs. Like), set liking, and decision certainty. Parametric regressors were not orthogonalized with respect to one another, allowing them to compete for variance independently. Missed trials (failures to choose a response within 5s in the Selection period) occurred rarely (0.4% of trials) and were modeled as a separate condition. As with our primary analyses, trials were concatenated and appropriate regressors were included to account for block-wise effects, and the GLMs were estimated with RobustWLS. We performed second-level random-effects analyses on the beta estimates generated at the first level, and whole-brain group statistical maps were generated using one-sample t-tests over these contrasts. These maps were extent-thresholded to achieve a whole-brain family-wise error cluster-corrected $p<0.05$. To avoid obscuring areas of potential overlap, we used a cluster-forming threshold of $p<0.005$ to generate these maps. However, despite the fact that these maps are confirmatory and therefore not used as the basis for statistical inference, in order to guard against potential false-positives[1,2] we separately confirmed that the same clusters remain significant with a more conservative cluster-forming threshold of $p<0.001$. These maps were projected onto the Caret-inflated cortical surface[3].

### Supplementary Results

*1. Alternate value difference formulation*

Our analyses estimate decision certainty on Choose trials based on the absolute difference between the value of the chosen item and the average of the remaining items. We considered whether a better estimate might obtain from the difference between the chosen item and the next-best item. While this alternate, Chosen-Versus-Next variable also correlates negatively with RT ($\beta = -0.19$, $t(26.0) = -7.3$, $p < 0.001$) and positively with choice confidence ($\beta = 0.14$, $t(26.0) = -6.9$, $p < 0.001$), these effects were overall smaller than those observed for the Chosen-Versus-Remaining variable. When entering both into the same regression, we find that only Chosen-Versus-Remaining continues to predict RT ($\beta = -0.27$, $t(37.2) = -8.1$, $p < 0.001$) and confidence ($\beta = 0.25$, $t(27.8) = 6.2$, $p < 0.001$) (Chosen-Versus-Next $|ts| < 2.0$, $p > 0.05$).

*2. Tests for other decision value signals in Dorsal and Ventral Value Networks*

To confirm the specificity of our results relating the Dorsal and Ventral Value Networks to Like and Choose respectively, we ran separate regressions to test whether either network demonstrated other value-related signals during either task or across tasks. Focusing on the appraisal task or collapsing across all trials, we did not find significant correlates of the overall set value, the value of the set's chosen option, or the difference between the chosen and unchosen options (Table S2). Focusing these analyses on Choose trials, we found positive correlates of chosen value and value difference in the Dorsal Value Network, but not the Ventral Value Network (Table S2). However, because chosen value and value difference are correlated ($r = 0.58$), we included them together in a single GLM and found that activity in the dorsal network on Choose trials is only significantly associated with value difference (consistent with the decision certainty analyses above; $\beta = 0.10$, $p < 0.05$) and not otherwise associated with the value of the chosen item ($p = 0.64$) or the overall set value ($p = 0.39$) (Table S4). None of these variables were significant predictors of activity in the Ventral Value Network (Table S4).

*3. Analysis of subjective choice confidence*

We also tested whether either network tracked subjective ratings of confidence for Choose decisions taken at the end of the experiment (during either task and across tasks), and did not find any such correlates ($|t|<1.1$, $p>0.30$). However, in addition to being retrospective, we note that these ratings were heavily skewed towards ceiling-level confidence ratings (mean = 4.25 [out of 5], SD = 1.06, median = 5). This was one of the motivations for focusing our decision certainty analyses on a continuous estimate of Choose certainty (value difference) that had a direct analog on Like trials (rating extremity), both of which having been measured at decision time.

### Supplementary Figures

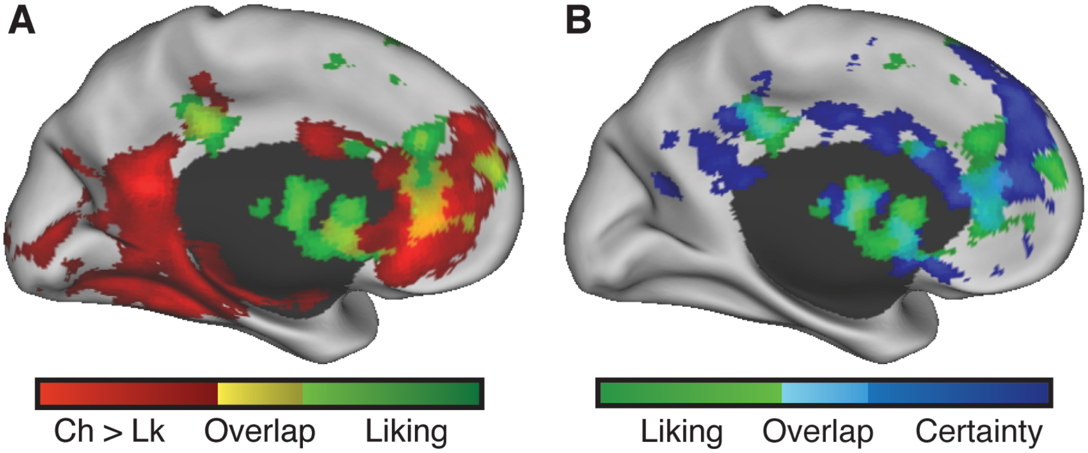

**Figure S1**. **A)** Confirmatory whole-brain analyses show that ventral striatum and more dorsal regions of vmPFC and PCC tracked set liking across tasks (green), whereas more ventral regions of vmPFC/PCC was significantly more active for Choose relative to Like trials (red), overlapping the set liking network (yellow). **B)** Decision certainty also activated regions of ventral striatum and dorsal regions of vmPFC/PCC (blue), overlapping regions associated with set liking (turquoise). Whole-brain statistical maps are thresholded at a cluster-corrected family-wise error (FWE) $p<0.05$.

### Supplementary Tables

| Choose > Like | | | | | |
|---|---|---|---|---|---|
| **Region** | **Cluster-level p-value** | **Cluster size (voxels)** | **Peak Voxel Z-score** | **Peak Voxel p-value** | **MNI Coordinates (mm)** |
| L Retrosplenial, parahippocampal cortex, calcarine, LOC | <0.00001 | 10051 | 5.98 | <0.00001 | -6,-56,18 |
| vmPFC, L MFG | <0.00001 | 4704 | 5.70 | <0.00001 | -6,38,0 |
| L Lateral OFC | 0.043 | 283 | 5.08 | <0.00001 | -34,32,-14 |
| R STG, operculum | 0.0018 | 498 | 4.87 | <0.00001 | 48,-8,-8 |
| R Calcarine, LOC | <0.00001 | 1308 | 4.81 | <0.00001 | 18,-78,12 |
| L Posterior insula | 0.036 | 294 | 4.29 | <0.00001 | -40,-4,-8 |
| B Cerebellum | 0.039 | 290 | 3.75 | 0.00009 | 2,-54,-36 |

| Like > Choose | | | | | |
|---|---|---|---|---|---|
| **Region** | **Cluster-level p-value** | **Cluster size (voxels)** | **Peak Voxel Z-score** | **Peak Voxel p-value** | **MNI Coordinates (mm)** |
| R DLPFC, IFG | <0.00001 | 3664 | 5.43 | <0.00001 | 46,38,22 |
| R IPL | <0.00001 | 1039 | 5.25 | <0.00001 | 50,-36,52 |
| L IFG | 0.0026 | 470 | 4.81 | <0.00001 | -48,14,6 |
| L IPL | <0.00001 | 954 | 4.69 | <0.00001 | -36,-44,44 |
| PCC | 0.023 | 322 | 4.38 | <0.00001 | -4,-24,28 |
| R MTG | 0.044 | 282 | 4.35 | <0.00001 | 58,-46,-8 |
| R SMA, SFG | 0.0053 | 421 | 3.75 | 0.00009 | 6,28,48 |

| Set liking (all trials) | | | | | |
|---|---|---|---|---|---|
| **Region** | **Cluster-level p-value** | **Cluster size (voxels)** | **Peak Voxel Z-score** | **Peak Voxel p-value** | **MNI Coordinates (mm)** |
| B VS/caudate, pgACC, L DLPFC, L MFG | <0.00001 | 7532 | 5.10 | <0.00001 | -34,6,58 |
| L MTG | 0.0000614 | 806 | 5.02 | <0.00001 | -58,-44,-16 |
| B PCC | 0.0164 | 362 | 4.40 | <0.00001 | 6,-32,36 |
| L IPL | 0.00012 | 750 | 4.05 | 0.00003 | -44,-46,44 |

| **Certainty (all trials)** | | | | | |
|---|---|---|---|---|---|
| **Region** | **Cluster-level p-value** | **Cluster size (voxels)** | **Peak Voxel Z-score** | **Peak Voxel p-value** | **MNI Coordinates (mm)** |
| B VS/caudate, pgACC, PCC, R IFG | <0.00001 | 10728 | 4.96 | <0.00001 | -4,46,18 |
| L MTG | <0.00001 | 2526 | 4.74 | <0.00001 | -56,-24,-12 |
| R amygdala | 0.0022 | 477 | 4.55 | <0.00001 | 26,-2,-18 |
| L precentral, IFG | <0.00001 | 1252 | 4.05 | 0.00003 | -10,-8,58 |
| R precentral | 0.033 | 297 | 3.85 | 0.00006 | 44,-8,58 |
| L postcentral | 0.016 | 343 | 3.64 | 0.00014 | -44,-28,44 |
| R MTG | 0.00013 | 691 | 3.62 | 0.00015 | 52,-54,14 |
| R TPJ | <0.00001 | 1026 | 3.58 | 0.00017 | 50,-8,22 |

**Table S1.** Results of whole-brain analysis for task (Choose vs. Like), set liking, and decision certainty. All of these were included in the same GLM, which also covaried RT. Whole-brain maps were thresholded at a voxelwise $p<0.005$, extent-thresholded to obtain a clusterwise $p<0.05$. LOC: lateral occipital cortex, MFG: middle frontal gyrus, STG: superior temporal gyrus, DLPFC: dorsolateral prefrontal cortex, IFG: inferior frontal gyrus, SFG: superior frontal gyrus, IPL: inferior parietal lobule, MTG: middle temporal gyrus, SMA: supplementary motor area, TPJ: temporoparietal junction.

| Network | Task | Predictor | β | SEM | t | p |
|---|---|---|---|---|---|---|
| **Dorsal Value Network** | **All trials** | Overall value | 0.04 | 0.02 | 1.68 | 0.106 |
| | | Chosen value | 0.04 | 0.02 | 1.91 | 0.067 |
| | | Value difference | 0.02 | 0.02 | 1.17 | 0.24 |
| | | \|Value difference\| | 0.03 | 0.02 | 1.59 | 0.12 |
| | **Appraisal** | Overall value | 0.03 | 0.03 | 0.98 | 0.335 |
| | | Chosen value | 0.00 | 0.03 | 0.10 | 0.919 |
| | | Value difference | -0.04 | 0.03 | -1.41 | 0.16 |
| | | \|Value difference\| | -0.04 | 0.02 | -1.75 | 0.08 |
| | **Choice** | Overall value | 0.05 | 0.03 | 1.75 | 0.092 |
| | | Chosen value | 0.08 | 0.03 | 3.06 | 0.005 |
| | | Value difference | 0.07 | 0.02 | 2.92 | 0.003 |
| | | \|Value difference\| | 0.10 | 0.03 | 3.84 | 0.0008 |
| **Ventral Value Network** | **All trials** | Overall value | 0.03 | 0.02 | 1.59 | 0.124 |
| | | Chosen value | 0.02 | 0.02 | 0.82 | 0.421 |
| | | Value difference | -0.01 | 0.02 | -0.84 | 0.403 |
| | | \|Value difference\| | 0.00 | 0.02 | -0.23 | 0.817 |
| | **Appraisal** | Overall value | 0.02 | 0.03 | 0.63 | 0.536 |
| | | Chosen value | 0.00 | 0.03 | 0.08 | 0.939 |
| | | Value difference | -0.02 | 0.02 | -0.71 | 0.48 |
| | | \|Value difference\| | -0.02 | 0.02 | -0.93 | 0.354 |
| | **Choice** | Overall value | 0.05 | 0.03 | 1.76 | 0.087 |
| | | Chosen value | 0.03 | 0.03 | 1.12 | 0.270 |
| | | Value difference | -0.01 | 0.03 | -0.53 | 0.60 |
| | | \|Value difference\| | 0.01 | 0.03 | 0.34 | 0.738 |

**Table S2**. Regression estimates for regressions predicting BOLD activity in the Ventral and Dorsal Value Networks based on overall value (average item value), chosen value (value of the item chosen from the set during the Choose task), and value difference (absolute distance between the chosen value and the average value of the remaining items). Each row represents a separate regression involving a single predictor. See the main text for relevant simultaneous regressions.

| **Outcome** | **Predictor** | **β** | **SEM** | **t** | **p** |
|---|---|---|---|---|---|
| **Set liking** | | | | | |
| | pgACC | 0.05 | 0.02 | 2.42 | 0.017 |
| | VS | 0.07 | 0.02 | 3.29 | <0.005 |
| | PCC | -0.02 | 0.02 | -0.72 | 0.47 |
| | Evaluation time | -0.05 | 0.02 | -2.03 | 0.053 |
| **Task condition** | | | | | |
| | mOFC | 0.16 | 0.06 | 2.72 | <0.01 |
| | MFG | 0.16 | 0.07 | 2.43 | 0.015 |
| | RSC | 0.17 | 0.05 | 3.09 | <0.005 |
| | Evaluation time | 0.24 | 0.07 | 3.32 | <0.001 |

**Table S3**. Regression estimates for regressions predicting set liking and task condition based on individual sub-regions within the Dorsal Value Network (pgACC, VS, PCC) and Ventral Value Network (mOFC, MFG, RSC), respectively, covarying evaluation time.

| **Outcome** | **Predictor** | **β** | **SEM** | **t** | **p** |
|---|---|---|---|---|---|
| **Dorsal Value Network** (Choose Trials) | | | | | |
| | Overall value | 0.04 | 0.05 | 0.86 | 0.39 |
| | Chosen value | 0.01 | 0.02 | 0.47 | 0.64 |
| | \|Value difference\| | 0.10 | 0.04 | 2.47 | 0.02 |
| | Evaluation time | 0.04 | 0.03 | 1.20 | 0.24 |
| **Ventral Value Network** (Choose Trials) | | | | | |
| | Overall value | 0.06 | 0.05 | 1.25 | 0.21 |
| | Chosen value | -0.00 | 0.02 | -0.06 | 0.95 |
| | \|Value difference\| | 0.04 | 0.04 | 0.87 | 0.39 |
| | Evaluation time | 0.09 | 0.03 | 3.12 | <0.005 |

**Table S4**. Regression estimates for Dorsal Value Network and Ventral Value Network activity on Choose trials, based on overall value, chosen value, value difference, and evaluation time.

| Outcome | Predictor | β | SEM | t | p |
|---|---|---|---|---|---|
| **Dorsal Value Network** | | | | | |
| | Condition | 0.10 | 0.04 | 2.57 | 0.011 |
| | Set liking | 0.08 | 0.03 | 2.93 | 0.007 |
| | Decision certainty | 0.05 | 0.02 | 2.35 | 0.020 |
| | Condition*liking | -0.03 | 0.04 | -0.74 | 0.46 |
| | Condition*certainty | 0.04 | 0.05 | 0.91 | 0.37 |
| | Evaluation time | 0.05 | 0.02 | 2.73 | 0.009 |
| **Ventral Value Network** | | | | | |
| | Condition | 0.15 | 0.04 | 3.86 | <0.001 |
| | Set liking | 0.07 | 0.03 | 2.51 | 0.016 |
| | Decision certainty | 0.01 | 0.02 | 0.60 | 0.55 |
| | Condition*liking | -0.01 | 0.04 | -0.24 | 0.81 |
| | Condition*certainty | 0.06 | 0.05 | 1.20 | 0.24 |
| | Evaluation time | 0.07 | 0.02 | 3.67 | <0.001 |

**Table S5**. Regression estimates for regressions predicting BOLD activity at the onset of the options (prior to task cue onset) in the Dorsal and Ventral Value Networks based on simultaneous predictors for task condition (Choose > Like), set liking, (task-relevant) decision certainty, condition x liking interaction, condition x certainty interaction, and (log) evaluation time.
